# Dissecting Epstein–Barr Virus Dependence Across Diverse Infected Cell Models

**DOI:** 10.64898/2025.12.19.695354

**Authors:** Alexandria Bartlett, Camille Krejdovsky, Guang Yang, Ashley P. Barry, Cliff Oduor, Jeffrey A. Bailey, Ann M. Moormann, Yoshitaka Sato, Micah A. Luftig

## Abstract

Epstein–Barr virus (EBV) is maintained clonally in a wide array of tumors and cells of lymphoproliferative disorders, yet the degree to which these infected cells continuously depend on EBV remains unresolved. To directly assess EBV dependence, we used two complementary Cas9-based viral eviction strategies: 1) targeting EBNA1 to promote episome dilution during cell division, and 2) targeting repetitive viral genomic regions to rapidly degrade EBV episomes. Using these approaches, we evaluated EBV-loss sensitivity across newly-derived, phenotypically diverse EBV-positive Burkitt lymphoma lines. We also evaluated cell lines from patients with EBV-associated T/NK and epithelial malignancies. We included the Burkitt Lymphoma cell line, Akata-GFP, as our EBV-loss-tolerant control and a lymphoblastoid cell line as our EBV-loss-sensitive control. We observed EBV-loss sensitivity in all the EBV-positive Burkitt Lymphoma cell lines and the chronic active EBV disease cell line. In contrast, the EBV-positive gastric adenocarcinoma lines were markedly less sensitive to EBV loss. EBV latency type did not predict EBV dependence, indicating that viral gene expression programs alone do not dictate reliance on the virus. These findings demonstrate that EBV dependence is prevalent across multiple EBV-associated disease models and is especially pronounced in early-state or newly derived lymphoid malignancies.

**AUTHOR SUMMARY:** Epstein–Barr virus (EBV) infects most people worldwide and is linked to many malignancies. Although EBV is present in cancer cells, it is unclear whether these cells require the virus to survive. To answer this question, we used CRISPR/Cas9-based methods to remove EBV from infected cells. Using these tools, we studied many different EBV-positive cell lines, including new Burkitt lymphoma cell lines. We found that most EBV-positive lymphoid cells required EBV for growth. In contrast, EBV-positive gastric cells were more tolerant of viral loss. These results show that many EBV-associated malignancies, especially early or newly established diseases, rely on EBV to survive.

## INTRODUCTION

Epstein-Barr virus (EBV) infects approximately 95% of adults worldwide and contributes to ∼1.5% of all human cancers [1]. Like other herpesviruses, EBV establishes life-long latent infections, but is unique in that it exists in a range of latency programs, characterized by varying profiles of viral gene expression [2]. In all latency programs, EBV Nuclear Antigen 1 (EBNA1) is expressed. EBNA1 is required for maintaining the multiple copies of the EBV episome as the infected cell divides [3].

EBV is clonally present in a wide array of tumor types and cells of lymphoproliferative disorders. The fact that the virus is maintained throughout proliferation of infected cells suggests that EBV plays a critical role in cell survival/proliferation. This has been investigated to some degree in the context of EBV-positive Burkitt Lymphoma (BL), the most prevalent pediatric cancer in sub-Saharan Africa. BL is defined by a c-myc translocation resulting in over-expression and ultimately uncontrolled tumor cell proliferation. There is currently no consensus regarding BL dependence on EBV.

Previous work has largely evaluated EBV dependence in BL through isolation of rare EBV-negative BL clones or through inhibition of EBNA1 via expression of dominant negative EBNA1 [4-6]. Studies comparing EBV-negative clones to EBV-positive parental lines have demonstrated EBV-negative clones to be more sensitive to apoptosis and less tumorigenic [5, 7]. We have expanded on these efforts to evaluate EBV dependence through the use of two different Cas9-based approaches to evict EBV in newly-derived BL cell lines. We targeted EBNA1 to dilute the EBV episome from infected cells as they divide [8]. We separately targeted repetitive regions of the EBV genome, to more rapidly degrade EBV episomes [9, 10]. In addition to evaluating EBV dependence in newly-derived BL cell lines, we also assessed EBV dependence in additional EBV disease models including chronic active EBV (CAEBV) and gastric adenocarcinoma cell lines.

## RESULTS/DISCUSSION

### Eviction in the EBV-loss-tolerant cell line, Akata-GFP

To evaluate EBV dependence, we aimed to evict EBV using two Cas9-based approaches. The first approach targeted EBNA1 (**Fig 1A**). EBNA1 is expressed in all latently infected cells and is responsible for maintaining the viral genome through cell division (**S1. Fig A**). Therefore, loss of EBNA1 is expected to result in viral genome loss as episome copy number is diluted through cell division. We used the Akata-GFP cell line to evaluate 5 single guide RNAs (gRNAs) targeting the 5’ coding sequence of EBNA1 (**S1. Fig B-C**). The Akata-GFP cell line was generated through infection of an EBV-negative Akata clone with a recombinant Akata virus that expresses eGFP [11]. Expression of eGFP thus serves as a proxy for EBV presence in this cell line. Transfection of the Akata-GFP cell line with EBNA1 gRNA5 resulted in the largest percentage of GFP-negative cells 14 days post transfection and was therefore used as the EBNA1 gRNA in future experiments.

**Figure 1.**
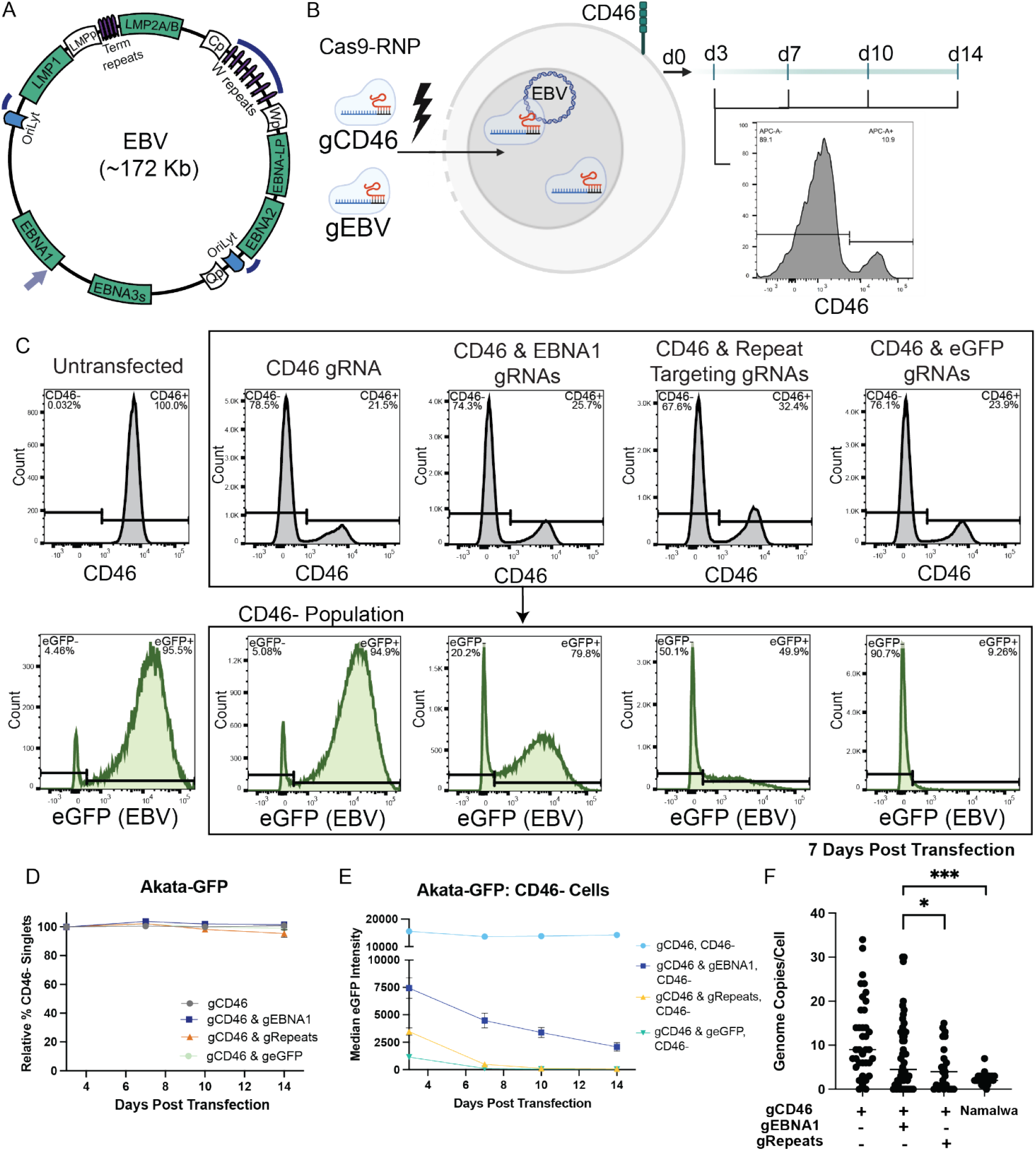
Cas9-RNP targeting of EBNA1 and EBV repetitive regions depletes EBV genomes, which is tolerated in Akata-GFP. **A)** EBV genome schematic with target sites of EBV directed gRNAs. EBNA1 gRNA indicated by lilac arrow and repeat targeting gRNAs indicated by navy curves. **B)** Cas9-RNP editing approach to evict EBV. EBV-positive cell lines are transfected with Cas9-RNPs targeting EBV (either EBNA1 or repetitive regions), and CD46. Cells are monitored for CD46 expression via flow cytometry starting at 3 days post transfection. Created in BioRender [15]. **C)** Representative flow cytometry histograms demonstrating CD46 and eGFP gating in Akata-GFP cells. Flow plots were taken from Day 7 Akata-GFP samples. **D)** Relative % CD46-negative cell populations of **Akata-GFP** (Latency I, EBV-loss-tolerant) cell line following transfection with Cas9 complexed with gRNAs targeting CD46, CD46 and EBNA1, CD46 and repetitive regions or CD46 and eGFP. Values were normalized to Day 3 transfection efficiency (mean ± SD). n=3 or 4 for all time points and treatments. **E)** Median eGFP fluorescence intensity of **CD46-negative** cell population in Akata-GFP cell line following transfection with Cas9 complexed with gRNAs targeting CD46, CD46 and EBNA1, CD46 and repetitive regions or CD46 and eGFP. **F)** EBV genomes/cell determined by EBV DNA FISH of Akata-GFP cell line at 7 days post transfection. Akata-GFP cells were sorted for the CD46-negative population following transfection. Median represented by line, Mann-Whitney test used to assess significance (*P< 0.05, ***P< 0.001).

The second EBV eviction approach used gRNAs targeting repetitive regions of the viral genome. We used 3 gRNAs, each with 7-25 potential target sites to rapidly degrade the viral genome (**Fig 1A**) [9, 10]. To employ these EBV eviction approaches, we transfected Akata-GFP cells with Cas9-Ribonucleoprotein complexes (Cas9-RNPs) targeting EBV (either EBNA1 or repetitive regions) and the non-essential cell surface marker, CD46 (**Fig 1B**). CD46 was used as a control, and proxy for successful gene targeting as our group and others have previously described [12, 13]. The CD46-negative population (i.e. the edited population) was monitored over time via flow cytometry starting at 3 days post infection, the time at which peak KO of CD46 is observed.

We observed minimal variation in the CD46-negative population following transfection of the Akata-GFP cell line with Cas9-RNPs targeting CD46, CD46 and EBNA1, CD46 and repetitive regions, or CD46 and eGFP (non-essential virus targeting control) (**Fig 1D**). This observation aligns with previous reports that Akata is tolerant to EBV loss [5, 6, 11]. When we targeted virally expressed eGFP, we observed strong loss of eGFP expression, particularly by the Day 7 time point (**Fig 1C, 1E**), suggesting high editing efficiency of the viral genome despite the approximately 47 EBV genome copies/cell present in the Akata-GFP cell line (**S1 Table**). We validated successful EBV targeting and EBV genome loss at the protein and DNA level. We evaluated eGFP expression in the CD46-negative population and observed a decrease in median eGFP intensity for the EBV and eGFP targeting treatments but not for cells transfected with only the CD46 targeting gRNA (**Fig 1C, 1E**). Furthermore, this trend of eGFP loss was not observed in the CD46-positive population (**S1. Fig E**). We also confirmed loss of EBNA1 in the CD46-negative population at the protein level via western blotting (**S1. Fig F**). To validate EBV genome loss at the DNA level, we sorted transfected cells for the CD46-negative populations and imaged EBV genomes in cells using a fluorescent *in situ* hybridization approach, OligoDNA-PAINT [14]. We observed a significant decrease of EBV genomes/cell for either EBV targeting treatment compared to the control group (only CD46 targeted), at Day 7 and 14 post transfection (**Fig 1F, S1. Fig G, H**).

### LCL and newly derived eBLs are sensitive to EBV loss

We applied the EBV eviction approaches to two additional control cell lines, a Lymphoblastoid Cell Line (LCL) which requires EBV for proliferation and the EBV-negative cell line, BJAB. We observed a reduction in the CD46-negative population when targeting EBV in LCL1391 (**Fig 2A**). The loss of the CD46-negative population was most pronounced for cells that were transfected with Cas9-RNPs targeting EBV repetitive regions, consistent with a rapid destruction of the viral genome. We did not observe a change in the CD46-negative population for BJAB when either EBV targeting approach was applied, confirming EBV specificity of the method (**Fig 2B**). As an additional control, we targeted a non-essential viral gene (BLLF1) to confirm that the loss of the CD46-negative population in LCLs was a result of viral genome loss and not only Cas9 cleavage of the viral genome **(S2. Fig D, M)**.

**Figure 2.**
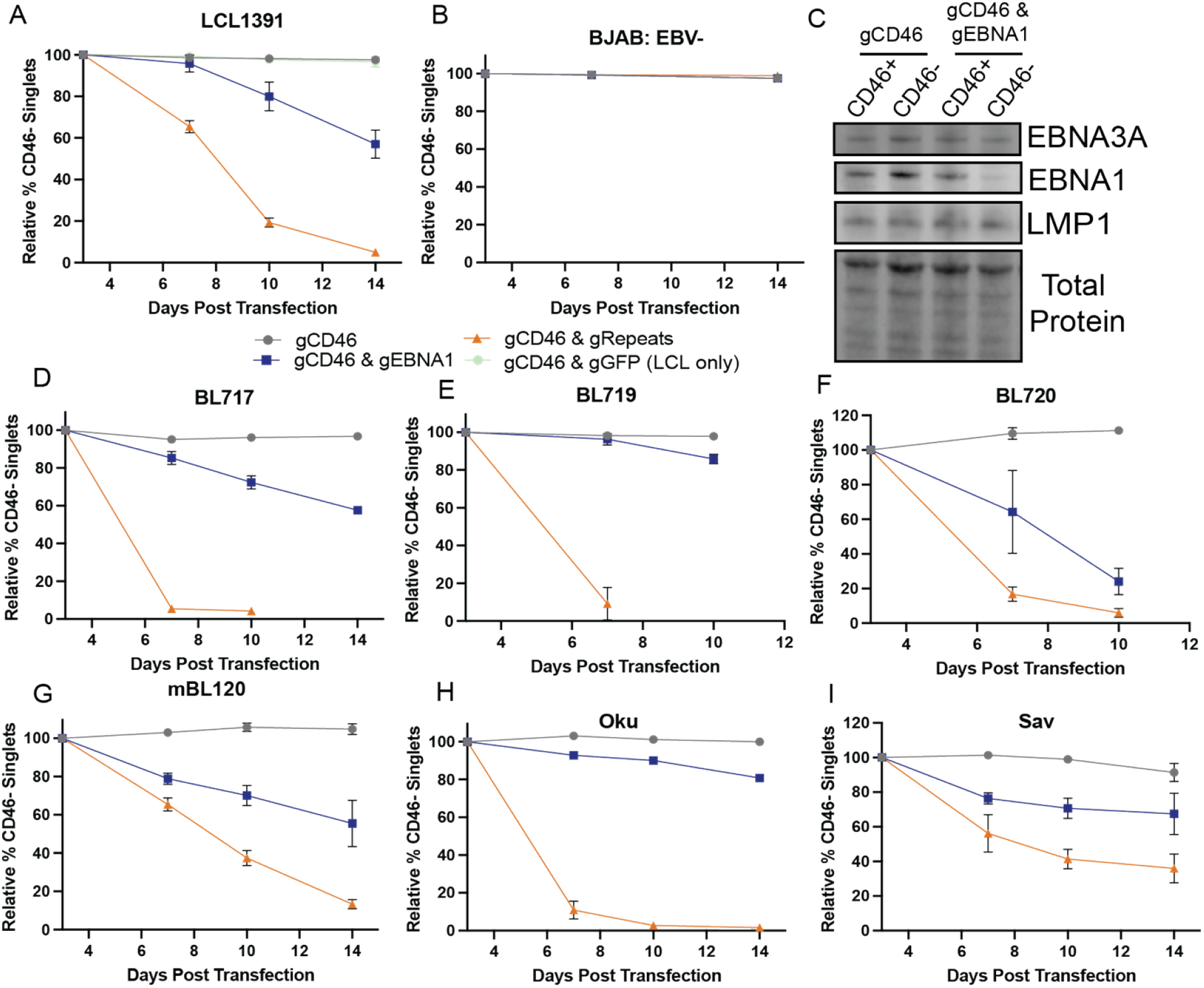
Cas9-RNP targeting of EBNA1 and EBV repetitive regions is not tolerated in LCL1391 or newly derived BL lines. **A)** Relative % CD46-negative cell populations of **LCL1391** (Latency III, EBV-loss-sensitive) cell line following transfection with Cas9 complexed with gRNAs targeting CD46, CD46 and EBNA1, CD46 and repetitive regions, or CD46 and eGFP (non-targeting control). **B)** Relative % CD46-negative cell populations of **BJAB** (EBV-negative) cell line following transfection with Cas9 complexed with gRNAs targeting CD46, CD46 and EBNA1 or CD46 and repetitive regions. **C)** Western blot of EBNA3A, EBNA1 and LMP1 expression in LCL1391 sorted on CD46 expression 3 days post transfection. Relative % CD46-negative cell populations following transfection with Cas9 complexed with gRNAs targeting CD46, CD46 and EBNA1 or CD46 and repetitive regions in **D)** BL717 (Latency IIb), **E)** BL719 (Latency III), **F)** BL720 (Latency III), **G)** MBL120 (Latency III), **H)** Oku (Wp-Restricted Latency), **I)** Sav (Latency I). For A,B, D-I, values were normalized to Day 3 transfection efficiency (mean ± SD). n=3 or 4.

At Day 3 post transfection of LCL1391, we confirmed decreased EBNA1 expression in the CD46-negative population using the EBNA1 targeting approach (**Fig 2C**). However, EBNA3A and LMP1 were still expressed, consistent with presence of the viral genome at this early time point and the slower viral genome dilution approach following EBNA1 loss. Furthermore, we confirmed editing of EBNA1 via Sanger sequencing and a KO score of 55% (**S2 Fig L**) [16].

We applied the EBV eviction approaches to four newly-derived BL cell lines (**S1 Table**). We have previously shown that the new BL cell lines represent diverse latency states, varying levels of spontaneous lytic activity and both type 1 and type 2 EBV strains [17, 18]. The newly derived BL cell lines were all sensitive to EBV loss regardless of latency state, lytic activity, strain or genome copy number (**Fig 2 D-G, S1 Table**). We found these results surprising given previous studies that were able to isolate EBV-negative clones from BL lines and observed BL clones that grew despite loss of EBV genomes through dnEBNA1 expression [4, 5]. Therefore, we also evaluated EBV dependence from BL lines in previous reports, Oku and Sav. The population loss of the CD46-negative population was more rapid with Oku compared to Sav for the repeats targeting method (**Fig 2H, I**). This observation is similar to a previously published study, in which one of two Sav clones was observed to proliferate despite EBV loss while both Oku clones were found to be sensitive to EBV loss [4]. However, ultimately our observations suggest Sav was still sensitive to EBV loss.

### EBV loss sensitivity varies in other disease models

To evaluate EBV loss sensitivity in other disease contexts, we applied the EBV eviction methods to an EBV-positive T cell line and two EBV-positive gastric adenocarcinoma cell lines. SNT-16 is an EBV-positive T cell line derived from a patient with CAEBV [19]. There was a progressive decrease in the CD46-negative population when we applied either EBV targeting approach to SNT-16 (**Fig 3A**). However, the gastric adenocarcinoma cell lines, SNU-719 and NCC-24, were markedly less sensitive to EBV eviction (**Fig 3B, C**). In particular, the CD46-negative population did not change when we applied either EBV eviction approach to NCC-24. It has previously been proposed that latency state may dictate dependence of EBV-positive cell lines on EBV [4]. EBV latency state varies between EBV-positive diseases (See Fig 2 in [2]). SNT-16, SNU-719 and NCC-24 cell lines represent similar latency states (latency IIa) yet have differing results regarding EBV-loss sensitivity. Therefore, our findings suggest viral gene expression program is not the main factor dictating dependence on EBV.

**Figure 3.**
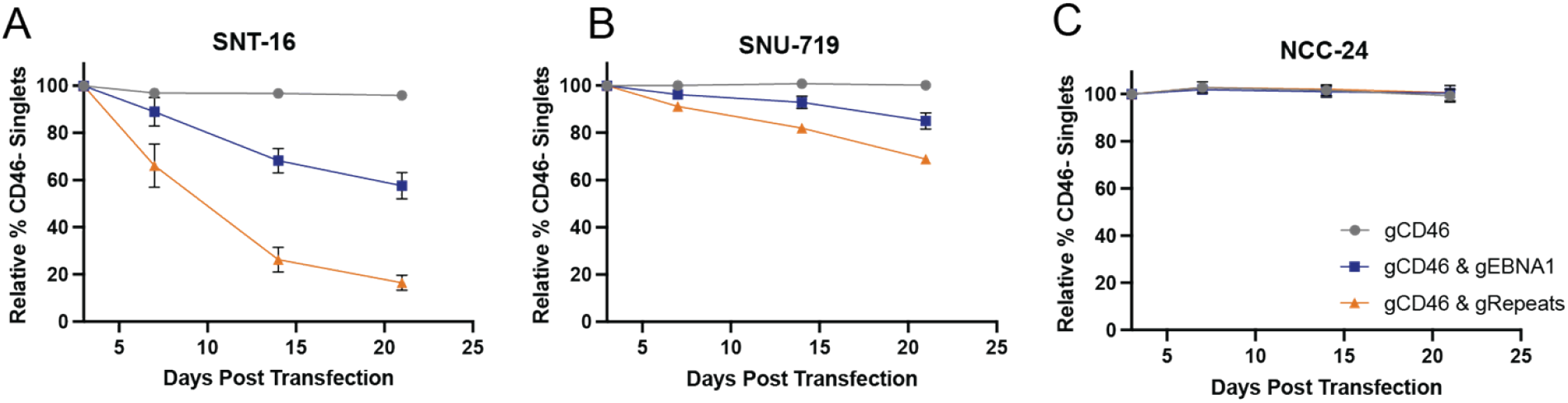
EBV dependence in other tumor types and lymphoproliferative disorders. Relative % CD46-negative cell populations following transfection with Cas9 complexed with gRNAs targeting CD46, CD46 and EBNA1 or CD46 and repetitive regions. **A)** SNT-16 (EBV-positive T cell line), **B)** SNU-719 (Gastric adenocarcinoma), and **C)** NCC-24 (Gastric adenocarcinoma).

## Conclusions

We have summarized the EBV loss sensitivity of the various cell lines evaluated in this study in Figure 4. EBV targeting of either EBNA1 or repetitive regions resulted in a decrease in the edited population for LCL1391 (EBV-loss-sensitive control), the BL lines and SNT-16. The gastric adenocarcinoma cell lines were markedly less sensitive to EBV loss, particularly NCC-24. There was a greater loss of the CD46-negative population in the EBV-loss-sensitive lines for the repetitive region targeting approach.

**Figure 4.**
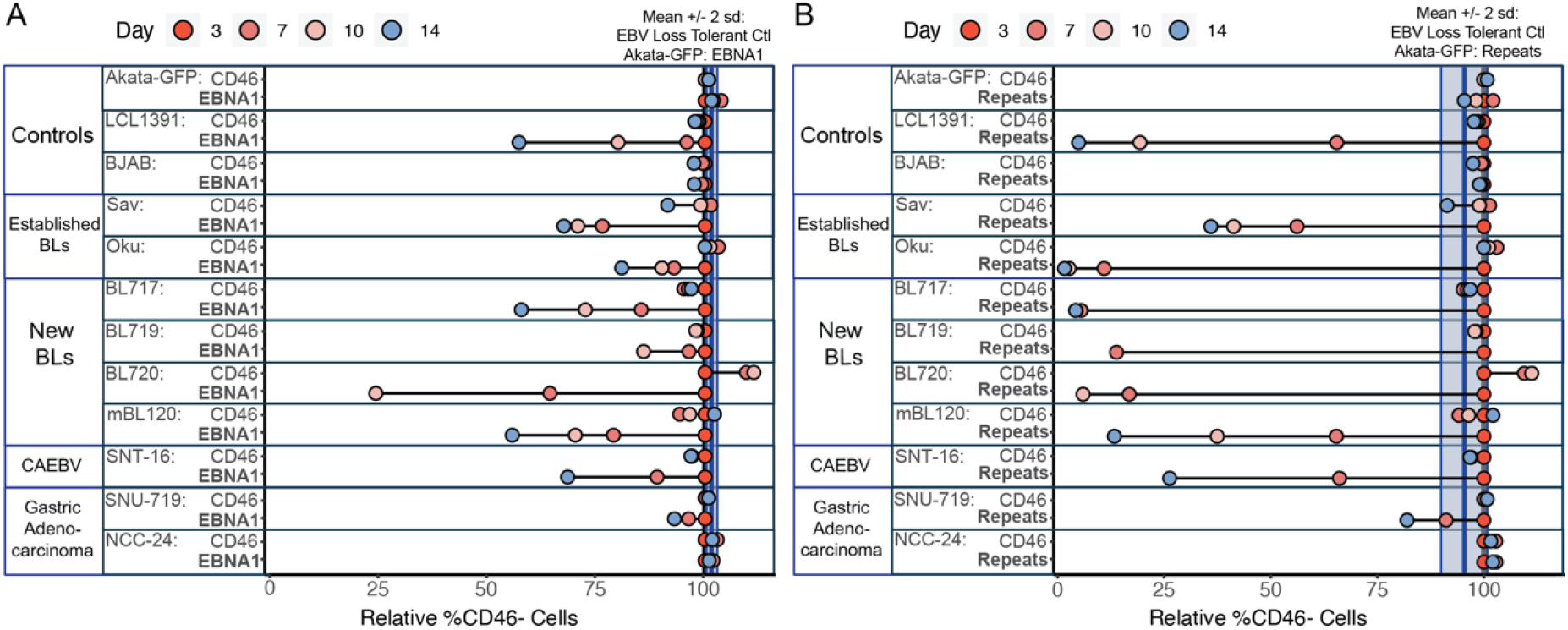
Summary of EBV-loss sensitivity in EBV-positive cell lines from different tumor and disease types. Relative % CD46-negative cell populations following transfection with Cas9 complexed with gRNAs targeting CD46, **A)** CD46 and EBNA or **B)** CD46 and repetitive regions. Light blue vertical bars represent the mean +/-2 sd for % CD46-negative cells at Day 14 for in Akata-GFP (EBV-loss-tolerant control) with EBV targeting (EBNA1 in A, Repeats in B). This is representative of the loss of the edited population in an EBV loss tolerant line. EBV targeting gRNAs resulted in a decrease in the edited population for LCL1391 (EBV-loss-sensitive control), and the majority of the EBV-positive cell lines profiled in this study.

Our findings provide support for a model in which EBV-loss sensitivity is prevalent in EBV-positive cell lines, including cell lines from different disease contexts and varying viral gene expression programs. The newly-derived BL and CAEBV cell lines were particularly sensitive to EBV loss, whereas the gastric adenocarcinoma cell lines were less sensitive to EBV loss. CAEBV often precedes EBV-positive T cell Lymphomas, therefore SNT-16 may represent an “early” state of the EBV-positive T cell disease spectrum. This is supported by fewer driver-gene mutations in CAEBV patients compared with other EBV-associated T/NK neoplasms [20]. Similar to the newly-derived BL lines, this early disease state may represent a time in disease when cells rely on the virus for proliferation. This concept is further supported by findings regarding the imperfect partitioning and maintenance of the viral genome by EBNA1, which would suggest loss of EBV if there was not selection [21]. However, cellular mutations may eventually allow an EBV-positive cell to evolve to be less reliant on the virus for proliferation [22]. Through this work EBV loss sensitivity emerges as a common feature of multiple EBV-positive tumor types, underscoring the continued selection for viral genome maintenance during proliferation of EBV-positive malignancies.

## Acknowledgments

We are greatly appreciative of Luftig laboratory members and our collaborators at The University of Massachusetts Chan Medical School, Brown University, The University of Zurich, and the Kenya Medical Research Institute for their insights and feedback. Flow cytometry analysis and sorting was conducted through the Duke Cancer Institute Flow Cytometry Core. This work was funded by NIH R01 CA234348 (to M.L., A.M., and J.B.) and NIH R01 CA189806 (to A.M.). A.B. was also supported through the Duke Center for Virology Viral Oncology Training Grant T32-CA009111.

